# Dendrite regeneration in *C. elegans*is controlled by the RAC GTPase CED-10 and the RhoGEF TIAM-1

**DOI:** 10.1101/2021.07.20.453023

**Authors:** Harjot Kaur Brar, Swagata Dey, Smriti Bhardwaj, Devashish Pande, Pallavi Singh, Shirshendu Dey, Anindya Ghosh-Roy

## Abstract

Neurons are vulnerable to physical insults which compromise the integrity of both dendrites and axons. Although several molecular pathways of axon regeneration are identified, our knowledge of dendrite regeneration is limited. To understand the mechanisms of dendrite regeneration, we used PVD neurons in *C. elegans* having stereotyped branched dendrites. Using femtosecond laser, we severed the primary dendrites and axon of this neuron. After severing the primary dendrites near the cell body, we observed sprouting of new branches from the proximal site within 6 hours, which regrew further with timein an unstereotyped manner.This was accompanied by reconnection between the proximal and distal dendrites as well as the fusion among the higher-order branches as reported before. We quantified the regeneration pattern in threeaspects –territory length, number of branchesand fusion phenomena.Axonal injury causes a retraction of the severed end followed by a Dual leucine zipper kinase-1 (DLK-1) dependent regrowth from the severed end.We tested the roles of the major axon regenerationsignaling hubs such as DLK-1-RPM-1, cAMP elevation, *let-7* miRNA, AKT-1, Phosphatidyl serine exposure/PS in dendrite regeneration. We found that neither regrowth nor fusionis affected by the axon injury pathway molecules. Surprisingly, we found that the RAC GTPase CED-10and its upstream GEF TIAM-1 play a cell-autonomous role in dendrite regeneration. Additionally, function of CED-10 in epidermal cell is critical for post-dendrotomy fusion phenomena. This work describes a novel regulatory mechanism of dendrite regeneration andprovides a framework for understanding the cellular mechanism of dendrite regeneration using PVD neuron as a model system.

## Introduction

Functional nervous system of an organism requires intact neuronal processes and synaptic connections for the proper transmission of electrical signals. A deficit in the structural integrity in the cognitive areas of the brain leads to the manifestation of neuropathologies(Dierssen and Ramakers, 2006, Dindot et al., 2008, Garey et al., 1998, Kaufmann and Moser, 2000). Due to their sensitivity towards excitatory and inhibitory inputs, dendrites are often the sites of neurotoxic damage leading to severe dendritic dystrophy such asformation of dendritic varicosities, loss of dendritic spines, mitochondrial swelling and dysfunction and disruption of microtubules(Mizielinska et al., 2009, Park et al., 1996, Hori and Carpenter, 1994). One or more of these hallmarks of dendrite damage have also been observed in focal stroke or anoxic depolarization(Risher et al., 2010), mild Traumatic Brain Injury (mTBI)(Gao et al., 2011) and epilepsy(Swann et al., 2000). Though these features may appear neuroprotective and reversible in favorable conditions, their frequent or chronic occurrence may be devastating or fatal. Unlike axonal damage and regeneration, dendrite regeneration has not been comprehensively explored.

The knowledge about neurite regeneration has been attainedmostly from the axonal injury models. An injury to the axons elicitsa local calcium increase(Gitler and Spira, 1998, Ghosh-Roy et al., 2010)that triggers elevation in the Cyclic Adenosine monophosphate (cAMP) levels, activation of downstream Protein Kinase A (PKA) and mitogen-activated protein kinase kinasekinase (MAPKKK), Dual Leucine Zipper Kinase (DLK-1) (Hammarlund et al., 2009, Yan et al., 2009, Hao et al., 2016). DLK-1 initiates local microtubule remodeling(Ghosh-Roy et al., 2012) and activatesEts-C/EBP-1 transcription complex promoting axon regeneration (Li et al., 2015).The Dendritic arborization (*da*) neurons in *Drosophila* have been recently established as an efficient model for studying dendrite regeneration(Song et al., 2012, Thompson-Peer et al., 2016). Both neuron intrinsic as well extrinsic machineries can regulate the efficiency of dendrite regeneration (DeVault et al., 2018). The dendrite regeneration is independent of Dual Leucine zipper Kinase (DLK) MAPK pathway (Stone et al., 2014), which is an essential factor for the initiation of axon regeneration (Hammarlund et al., 2009).However, other kinases like AKT, and Ror have been implicated in the process(Song et al., 2012, Nye et al., 2020). Also,Wnt effectors, which regulate the dendritic morphology and branching can also regulate the process of dendrite regeneration(Weiner et al., 2020, Nye et al., 2020).Although some of the cytoskeleton-based mechanisms controlling the axon regrowth do not affect dendrite regeneration(Rao et al., 2016), microtubule minus end binding protein, Patronin-1 controlsboth axon and dendrite regeneration(Feng et al., 2019, Hertzler et al., 2020, Chuang et al., 2014). The roles of the axon regeneration machineries have not been extensively tested for dendrite regeneration.

PVD neuron in *C. elegans*which is responsible for harsh touch sensation, hasan elaborate dendritic branching pattern (Chatzigeorgiou et al., 2010, Tao et al., 2019). Laser induced small damage to the dendrites of PVD neuron triggers a regenerative self-fusion process (Oren-Suissa et al., 2017, Kravtsov et al., 2017). The fusogen AFF-1 plays an important role in promoting the fusion between the injured proximal and distal dendrites (Oren-Suissa et al., 2017). However, the early signaling mechanisms leading to initiation of dendrite regrowth remain elusive.

In this report, by combining 2-photon laser neurosurgery and quantitative imaging, we have characterized the PVD neuron system in worms for both axon and dendrite injury paradigms. Using both dendrite and the axon regeneration assays in the same neuron, we assessed the roles of axon regeneration pathways in dendrite regeneration.Our results showed that the dendrite regeneration is a multivariate process comprising of regrowth, branching and fusion events, which are independent of conventional axon regeneration pathways including DLK-1/MLK-1. Our results highlight the neuronal and epidermal roles of RacGTPase, CED-10 in the initiation of dendrite regrowth and self-fusion processes. We also showed that theTIAM-1, aRho Guanine Exchange Factor (Rho GEF)acts upstream to CED-10 for dendrite regrowth and branching.

## Results

### Primary dendrite injury in PVD neuron triggers multiple responses involving regrowth, branching, and fusion

In *C. elegans*, PVD neurons are located mediolaterally with a well-defined ventrally targeted axon and dendritic structure that spans in anterior-posterior direction and their orthogonal arbors reaching the dorsal and ventral midline (Figure 1A). These dendrites are hierarchically classified from primary to quaternary based on theirorderof branching(Oren-Suissa et al., 2010). Previous studies have elucidated that following injury, primary major dendrites of PVD neurons sprout neurites, which quickly fuse with their distal counterparts with the help of the fusogen, AFF-1 delivered from the epidermal cells(Oren-Suissa et al., 2017). The mechanisms that initiate regeneration process after dendrotomyare yet to be investigated. Using GFP and mCherry∷RAB-3 labeled PVD neurons (Figure 1A), we identified the axonal and primary dendritic compartments of PVD and performed dendrotomy with a modified paradigm (Figure 1A).We delayed the self-fusion process by creating abig gap between the proximal and distal part of primary dendrite by using two successive laser-shots at 10-15μm apart (Red arrow, Figure 1B). The dendrites regenerated from the severed end as well as from the nearest proximal tertiary dendrites (3h post-dendrotomy),subsequently branched more(6h post-dendrotomy) (Figure 1B) and eventually reconnectedtothe distal dendrites at 12h and 24h post injury (green arrowheads,Figure 1B,). This process was also accompanied by menorah-menorah fusion(Oren-Suissa et al., 2017), in which the tertiary dendrites adjacent to the injury site merge with each other to bypass the gap of injury(red semi-transparent box,Figure 1B) and degeneration of the distal part (Figure 1B, gray traces in schematics). As time progressed (48h), the regrowing dendriteexpaned the territory furtherwith an increased number of branches(Figure 1B). Thelongest regrowing neurite (yellow dotted trace, Figure 1B)was used to quantitate the territory covered by these regenerated neurites, termed as ‘territory length’.The territory length increased with respect to time after dendrotomy (Figure 1C).The regenerative branching also showed an increase in number with time after dendrotomy (Figure 1D).The other parameters such as ‘reconnection’ with the distal dendrite and ‘menorah-menorah fusion’ as described earlier(Oren-Suissa et al., 2017), also increased with time after injury(Figure 1E-F). As we have seen that after dendrotomy, both regrowth and fusion happen simultaneously, we asked whether the fusion process couldprevent the extent of dendrite regeneration.

**Figure1.**
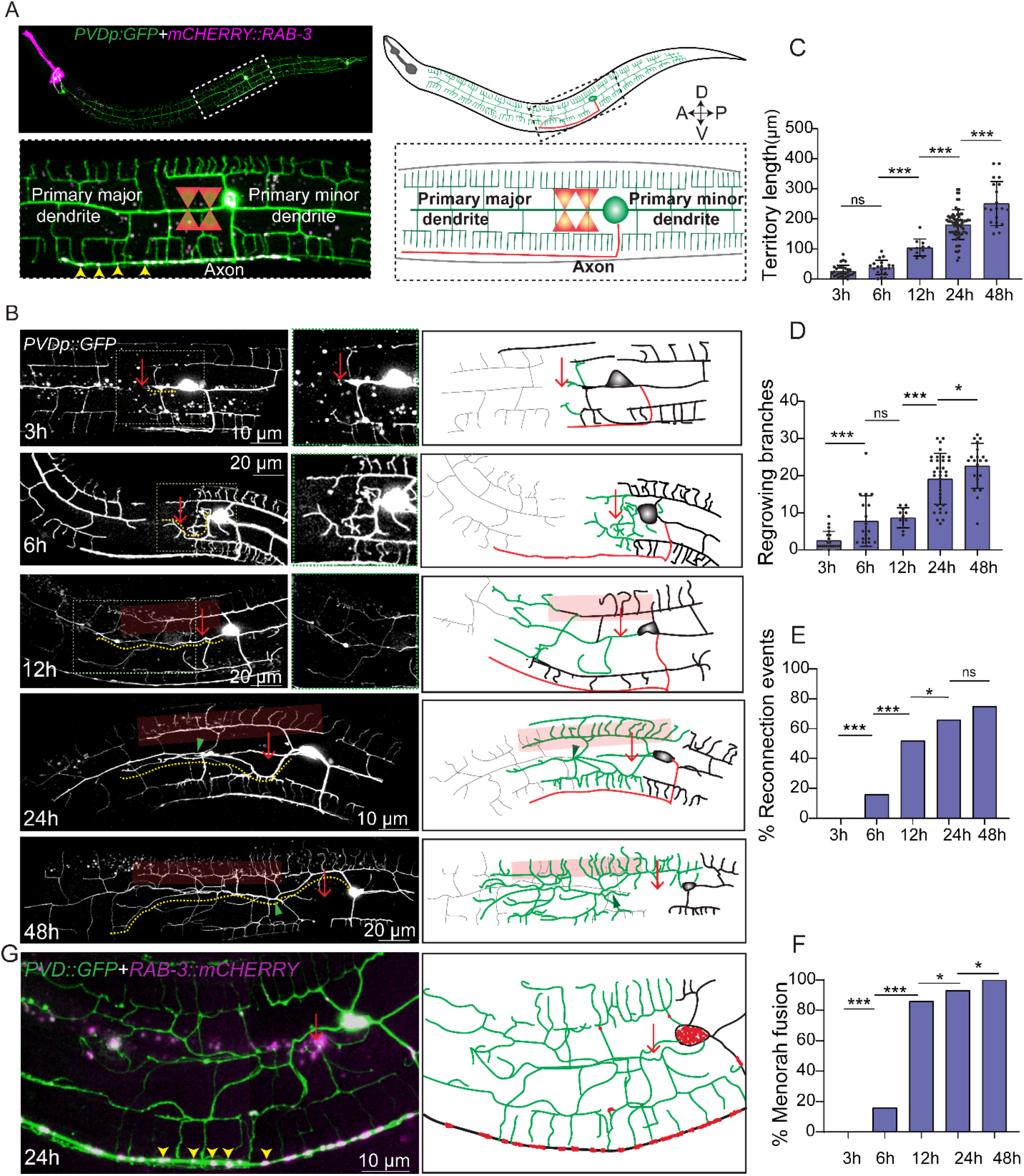
Primary dendrites of PVD neuron show multiple regenerative responses followingdendrotomy. (A) Representative confocal image and illustration of PVD neuron labeled with soluble GFP*wdIs52* (*pF49H12.4∷GFP*) and axon marked with mCherry∷RAB-3 punctae (*kyIs445*,P*des-2∷mcherry∷RAB-3*). The white dashed boxis magnified below and the primary major, primary minor dendrites and the axon are clearly highlighted.The yellow arrowheads are marking the axonal RAB-3 punctae. Schematics indicate dendrites in green and axon in red.The two shot dendrotomy cut using femtosecond laser is illustrated onto the magnified image and schematics. (B) Confocal images and schematics (right) showing the PVD neuronsdendrotomized with two laser shots in wild-type background at 3h, 6h,12h, 24h and 48h after injury. The PVD neuron is expressing the *pF49H12.4∷GFP*(*wdIs52)* reporter. At 3h after injury, the gap caused due to the two laser shots is shown with red arrow. A magnified view of the regrowing end within the green dotted box is shown on the right. The faded red boxes highlight menorah-menorah fusion events and the green arrowhead represents reconnection events between the proximal and distal primary dendrites. In the schematic views of the regeneration events, regenerated part and the distal remnants are shown in green and grey colors, respectively whereas the axon is shown in red. The longest regrowing dendrite is indicated with a yellow dotted line. (C) Quantification of longest regrowing dendrite, which is referred as ‘territory length’, (20≤n (number of regrowth events)≤70,N (Independent replicates)≥3), (D) Number of regrowing branches (14≤n≤34,N≥3) in each timepoint. compared using one-way ANOVA with Tukeymultiple comparisons method. Data is represented as a column scatter and histogram of Mean±S.D. p<0.05*, 0.01**, 0.001***, ns (non-significant, p>0.05). (E-F) The percentage occurrence of reconnection events (20≤n≤60,N≥3), and the menorah-menorah fusion (20≤n≤60,N≥3). The Fisher exact two-tailed test was done for statistical analysis. ns, non-significant, p<0.05*, 0.01**, 0.001***. (G) Confocal images of dendrotomized PVD neuron showing the localization of mCherry∷RAB-3 labeled with GFP (*F49H12.4∷GFP*)*wdIs52* and RAB-3 punctae with mcherry (*pdes-2∷mCherry∷rab-3*)*kyIs445* at 24h after dendrotomy with schematics showing localization of RAB-3 punctaeusingyellow arrowheads in confocal image and red dots in schematics. Regenerated dendrites are labeled as green traces.

To delay the fusion process further,we severed the dendrites usingfour consecutive shots which created abigger gap ~100μm (Figure S1A, right panel).The gap created between proximal and distal dendritic part due to multiple shots was significantly biggeras observed at 3-6h after injury (orangedouble-headed arrow, Figure S1A).Although the menorah-menorah fusion and reconnectionevents were significantly lower in multi-shot experiments(Figure S1B-C), both the territory length and % branching in this experiment were comparable to that of two shot dendrotomy experiment (Figure S1D-E).This indicated the fusion processdoes not influence the regenerative growth initiated upon dendrotomy in PVD neurons. However, dendrite regeneration in PVD neurons appeared to be a multivariate process.The synaptic reporter RAB-3∷mCherry mostly remained at the ventral cord (yellow arrowheads, Figure 1G) and did not invade into the regenerated neurites after dendrotomy. Additionally, we haveperformed dendrotomy on minor dendrite (Figure S1F).We observed similar regrowth response and fusion events in minordendrites as well following dendrotomy. (Figure S1F).

Hence, both major and minor dendrites of PVD neuron are able to regenerate after dendrotomyirrespective of the size of the injury. The regrowing dendrites cover up the injury area in a pattern different from the original arbor. In the event of encounter with the distal remnants, the regrowing dendrites may fuse and integrate into the original arbor. This however, does not prevent the unfused dendritic tips to grow further.

### Dendrite regeneration in PVD neurons is independent of DLK/MLK pathway

The cellular and molecular mechanism of axon regeneration has been extensively studied using various model organisms (He and Jin, 2016).The conserved mitogen-activated protein kinase kinasekinase (MAPKKK) pathway involving Dual Leucine Zipper Kinase (DLK-1) is essential for the initiation of regrowth from the cut stump of axon in multiple model organisms including mammals (Hammarlund et al., 2009, Yan et al., 2009, Shin et al., 2012), (Figure 2A).Therefore, it is possible that initiation of dendrite regrowth might rely on DLK-1 pathway.

**Figure 2:**
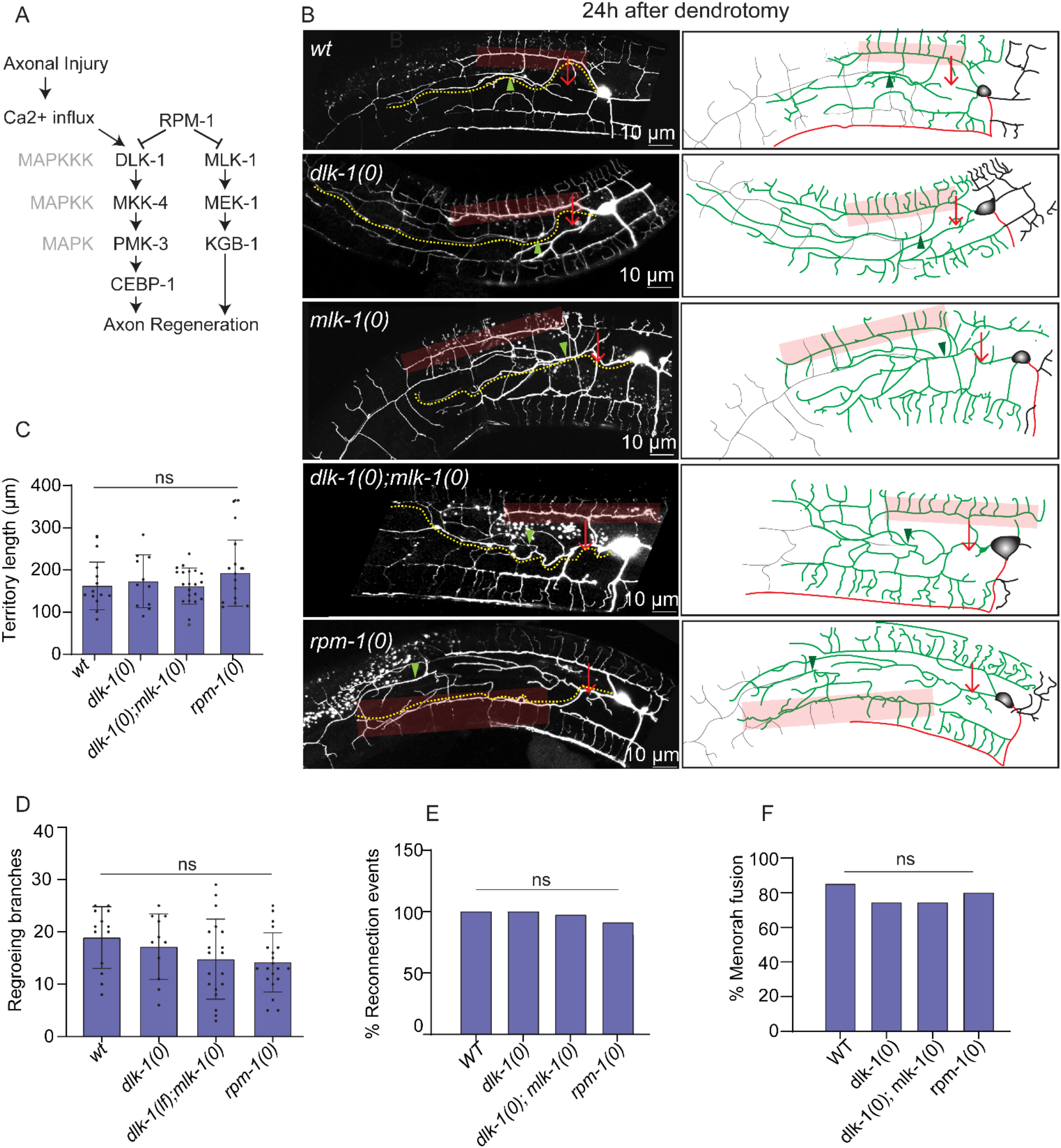
The dendrite regeneration is independent of DLK/MLK pathways. (A) Signaling pathway involving DLK-1 MAP kinase responsible for the initiation of axonal regeneration following axotomy. (B) Confocal images of the regeneration events of primary major dendrites in the wildtype, *dlk-1(0), mlk-1(0), dlk-1(0); mlk-1(0)* and*rpm-1(0)* at 24h post-dendrotomy. The experiment was done in the in *wdIs52*(*pF49H12.4∷GFP*) reporter background.The illustrations on the right indicating site of dendritic injury (red arrow), regenerating dendrites in green color, distal part in grey color, territory length in yellow dotted lines, fusion phenomena with green arrowheads and menorah-menorah fusion with faded red boxes. (C-F) Quantification of the territory length (C), total number of branches (D), the percentage of reconnection events (E), and the percentage of menorah-menorah fusionevents (F) in thewildtype and loss of function mutants of *dlk-1(0)*, *dlk-1(0);mlk-1(0)* and *rpm-1(0)* as (15≤n≤20, N≥3) at 24h post-dendrotomy. For C-D, statistics,One-way ANOVA with Tukeymultiple comparison test was done andfor E-F, the statistical comparison was done using Fisher exact two-tailed test considering ns,non-significant when p>0.05 and p<0.05*, 0.01**, 0.001***.

At 24-h following dendrotomy, the primary major dendrite in *dlk-1(0)*regrew in a manner similar to wild type (Figure 2B). The regenerative branching in the mutant was accompanied by reconnection of the primary dendrites (green arrowheads) and fusion between the tertiary dendrites equivalent to wild-type (red transparent box) (Figure 2B). Since both DLK-1 and MLK-1 cooperate to activate the regeneration response (Nix et al., 2011), we tested the single mutant *mlk-1(0)* as well as the double mutant lacking both*dlk-1* and *mlk-1.*In *mlk-1(0)* and *dlk-1(0); mlk-1(0),* dendrite regeneration was unaffected (Figure 2B). The quantitative parameters like territory length, number of regrowing branches in *dlk-1(0), mlk-1(0)* and *dlk-1(0);mlk-1(0)* were comparable to thewild-type (Figure 2B-D).Similarly, the reconnectionphenomena and menorah-menorah fusion events were equivalent in these mutants as compared to wild type (Figure 2E-F). The dendrite regeneration was also not influenced in the loss of function mutant of E3 ubiquitin ligase, RPM-1 (Figure 2B-F), which is known to downregulate DLK-1 and downstream kinases in the cascade during developmental growth of axon (Nakata et al., 2005). The dendrite regrowth and its ability to fuse at 24h after injury in *rpm-1(0)* was similar to the wild type (Figure2B-F) suggesting that *dlk-1* and *mlk-1* are neither necessary nor sufficient for the dendrite regrowth following injury in PVD neurons.

Furthermore, we checked the dendrite regeneration in the minor dendrite of *dlk-1(0);mlk-1(0)* double mutant, which was comparable to the wild-type (Figure S2A-C). These observations corroborate earlier results in *Drosophilada* neurons where dendrite regeneration was independent of the DLK-1signaling(Stone et al., 2014).Although*dlk-1* is expressed in PVD(Smith et al., 2010), its role in PVD neuron is unclear. Since a well-known role of E3 Ubiquitin ligase RPM-1 and downstream MAPKKK DLK-1 is to stabilize synaptic growth along with axon growth during development(Figure S2D)(Nakata et al., 2005, DiAntonio et al., 2001, Lewcock et al., 2007, Schaefer et al., 2000), we looked at the possible phenotype in axon development in *rpm-1* mutant.Both the *ju23* and *ok364* alleles of *rpm-1* showed an overgrowth of axon along the ventral cord (Figure S2E).The length of the axon is significantly higher in *rpm-1* mutants (Figure S2F) and axon overgrowth phenotype is completely suppressed by loss of *dlk-1* in *rpm-1(0)* background (Figure S2E-F) as seen in other neurons in *C. elegans*(Nakata et al., 2005)and other organisms(DiAntonio et al., 2001, Lewcock et al., 2007). This indicated that *rpm-1/dlk-1* cascade is functional in PVD neuron and strengthened our observation of unaffected dendrite regeneration in *dlk-1/mlk-1*mutants.

### Axon regeneration in PVD neurons depends on DLK-1 and MLK-1

Our finding that dendrite regeneration in PVD is independent of DLK-1 cascade raises a question whether the axon regeneration in this neuron would require this MAP Kinase pathway. We performed axotomy at 50 μmaway from the soma (Red arrow, Figure 3A) and found that at 3h after post-axotomy, the severed end retracted followed by a regrowth from the severed end afterwards (Figure 3B-C). The punctae of axonal reporter mCherry∷RAB-3 were localized at the tip of this regrowing neurite (Figure 3B, yellow arrowheads). These punctaeare oftenrelocalizedat the adjacent dendrites (Figure 3B, yellow arrowheads) suggesting conversion of someof the adjacent tertiary dendritesinto an axon. This observation was reminiscent of *Drosophilada* neurons where the dendrites are converted to axon following a proximal axotomy(Stone et al., 2014). The proximal part of the severed axon and converted neurites emanated some ectopic processes (Figure 3B, orange arrowheads)which too had mCherry∷RAB-3punctae (Figure3B). There was a significantextensionof the axon from the severed end at 24h and 48h as compared to 3h post-axotomy (Figure3B-C). Similarly, there was an increase in the conversion of dendritic branches into axon (Figure 3C) and the number and length of ectopic branches (Figure 3C).

**Figure 3:**
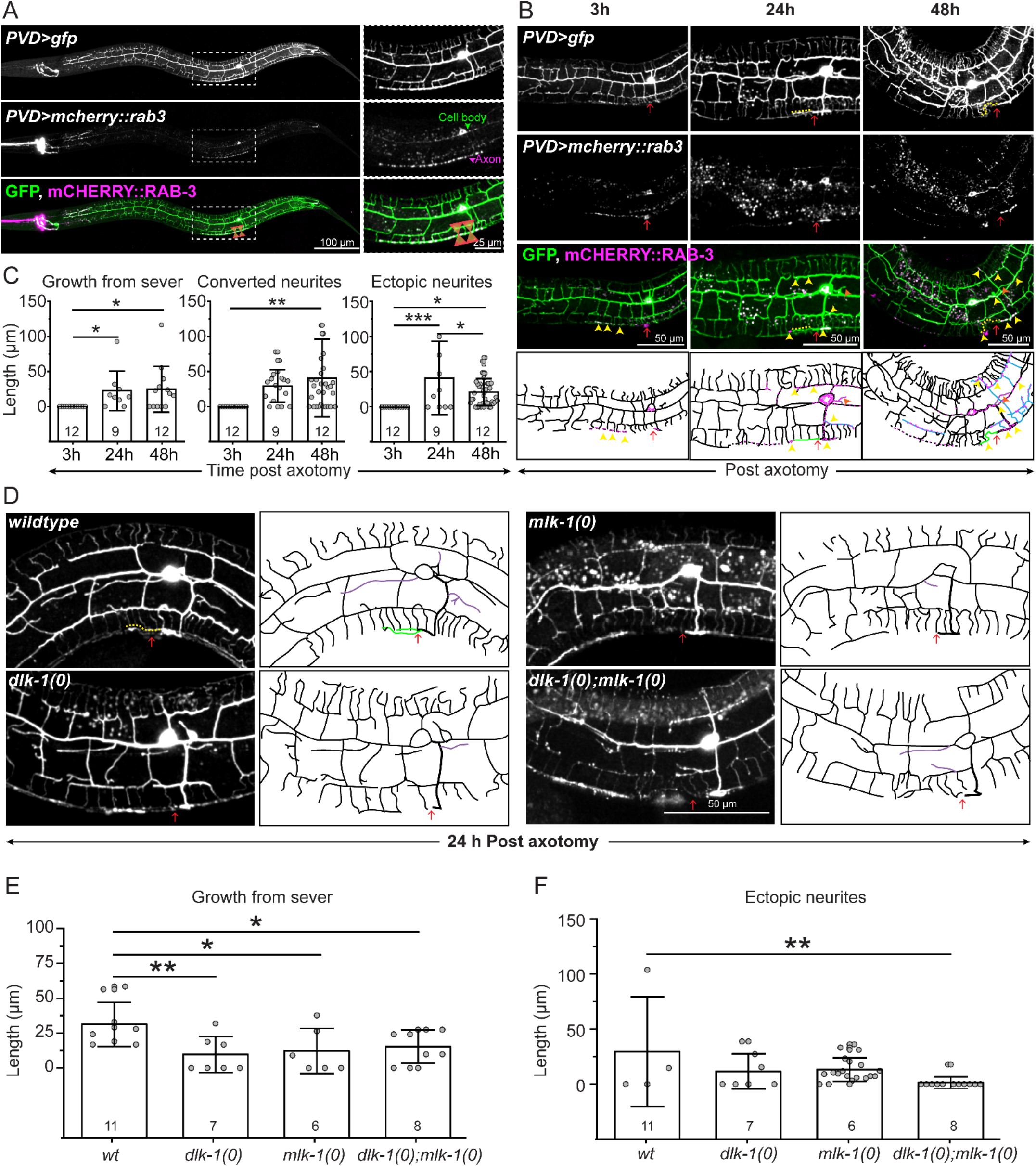
Axon regeneration in PVD neuron requires the DLK-1/MLK-1 pathway. (A) Representative images of PVD neuron labeled with *wdIs52* (*pF49H12.4>gfp*, green) and *kyIs445* (*pdes-2>mCherry∷rab-3*, magenta) with the region of interest (white dashed rectangle) magnified in the respective insets. mCherry∷RAB-3 punctae localized to the cell body and axon (marked in the inset). The axotomy injury using femtosecond laser is illustrated in the merged image. (B) Representative images and schematics at 3, 24, and 48h after axotomy of L4 stage (red arrow) PVD neurons labeled with GFP and mCherry∷RAB-3 (*kyIs445;wdIs52*) in green, magenta and merge channels. Based on the relocalization of mCherry∷RAB-3 (magentadots in schematics) (yellow arrowheads), the schematics show regenerative growth from the severed end (green traces), converted neurites (blue traces), ectopic neurites (purple traces in schematics)(orange arrowheads) classified as axonal branches. Axonal injury is marked using red arrow. (C) Quantification of the axon regeneration at 3, 24, and 48h following axotomy in the form of growth from the severed end, length of the converted neurites, and ectopic neuritesand compared using ANOVA, p<0.05*, 0.01**, 0.001*** (9≤n≤12,N≥3). (D) Regenerative growth after 24h of axotomy of the GFP labeled L4 stage PVD neurons in wildtype, *dlk-1(0)*, *mlk-1(0)*, and *dlk-1(0);mlk-1(0)* mutants with their representative images and schematics. The schematics show regenerative growth from the severed end (green traces) as axonal branches and ectopic neurites as purple traces. Axonal injury is labeled with red arrow.(E-F) Axon regeneration is quantified as growth from the severed end (E), and ectopic neurites (F) (6≥n≥11,N≥3) using ANOVA, p<0.05*, 0.01**, 0.001***.

We then carried out the axotomy in loss of function mutants of *dlk-1* and *mlk-1* (Figure 3D). At 24h post-axotomy, wildtype worms showed an average regrowth of 31.3±15.8 μm from the severed end which decreased significantly, due to loss of either *dlk-1* (9.7±12.9 μm) or *mlk-1* (12.2±16.2 μm) or both (15.4±11.8 μm) (Figure 3D-E), with negligible regrowth in more than 50% of the mutant worms. Length of the ectopic branches formed during regrowth also tended to decrease due to loss of *dlk-1* or *mlk-1* but double mutant showed a significant decrease in the same (Figure 3F). This confirmed the requirement of *dlk-1* and *mlk-1* in the PVD axon regeneration.

Thus the axon regeneration requires DLK/MLK pathway in PVD neuron but the dendrite regeneration is not dependent upon this signaling pathway as also seen in *Drosophila*(Stone et al., 2014). However, dendrite regeneration might rely on other molecular pathways regulating axon regeneration.

### Dendrite regeneration in PVD neurons is independent of conventional axon regeneration pathways

Axon regeneration is also controlled by pathways other than the DLK-1 pathway(Bradke et al., 2012). We tested some of the major genetic regulators implicated in axon regrowth.Axon regeneration is controlled by conserved Calcium and cAMP cascade in many organisms(Qiu et al., 2002, Ghosh-Roy et al., 2010). After axonal injury, there is a calcium influx (Gitler and Spira, 1998, Ghosh-Roy et al., 2010),which triggers acAMP cascadenear the injury site andactivates DLK-1 MAP3K(Hao et al., 2016). An elevation of either intracellular calcium using a gain of function mutation in L-type voltage gated calcium channel *egl-19*or an elevation of cAMP due to the loss of neuronal phosphodiesterase *pde-4* promotes axon regeneration (Ghosh-Roy et al., 2010). However, we observed that neither *egl-19(gf)*nor *pde-4(lf)* influenced any aspect of dendrite regeneration (Figure 4A-C).After 24hours of dendrotomy, the dendrite was able to regenerate to similar extent as wild type and the fusion phenomena werealso similar to wild-type (Figure 3 A-C).

**Figure 4.**
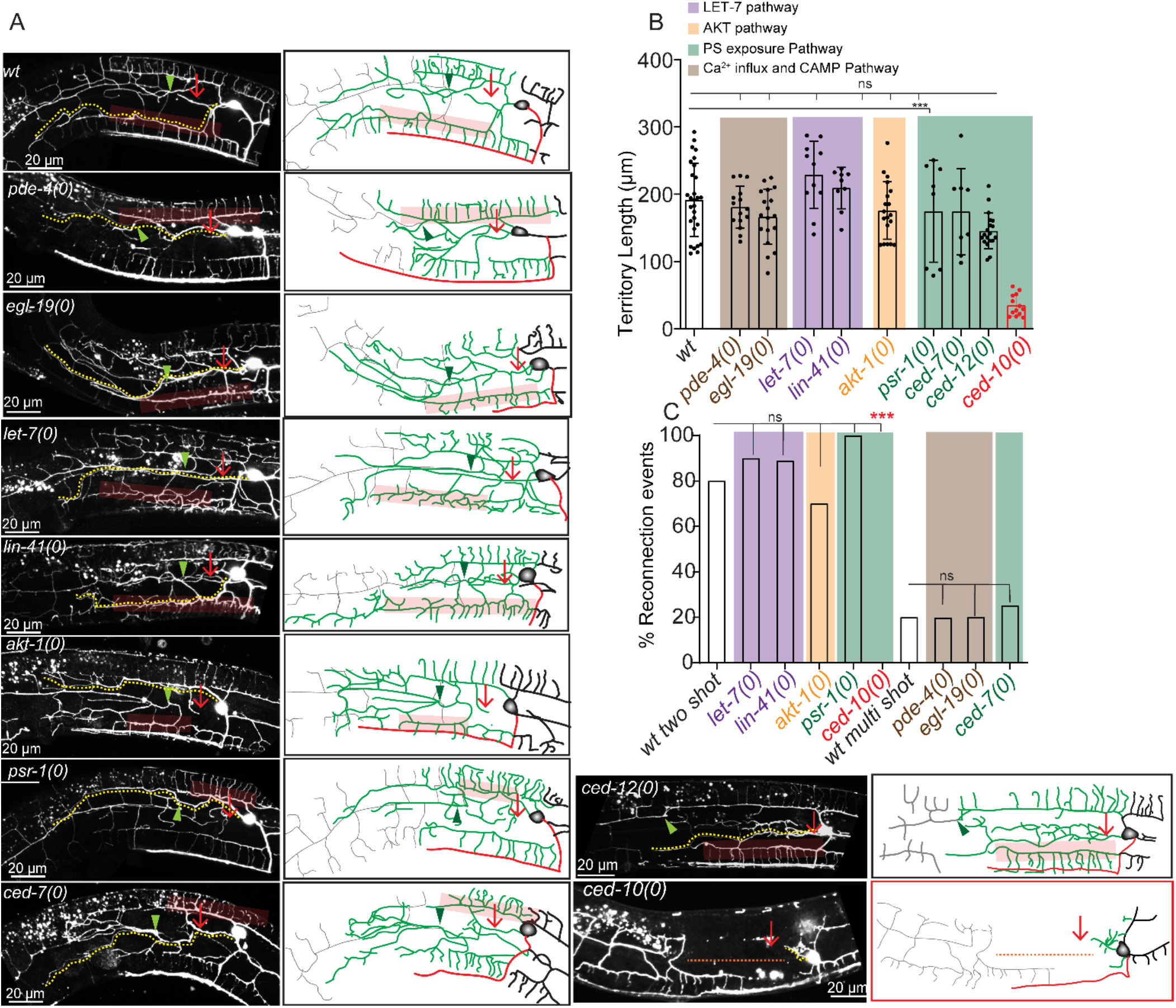
Dendrite regeneration is independent of conventional axon regeneration molecules. (A) Confocal images of the major dendrite regeneration events in wild type background, *pde-4(0), egl-19(0), let-7(0), lin-41(0), akt-1(0), psr-1(0), ced-7(0),ced-12(0), and ced-10(0).* The experiment is conductedusing*wdIs52* (*pF49H12.4*∷GFP) reporter.The illustrationson the right indicatingsite of dendritic injury (red arrow), regenerated dendrites (green), distal dendritic part (grey), territory length (yellow dotted lines), fusion like phenomena (green arrowheads) and menorah-menorah fusion (faded red boxes).(B-C) The quantification of the territorylength (B) and the percentage of reconnection events (C) obtained from the dendrotomy experiments shown in A (8≤n≤19, N≥3). Statistics, For B, One-way ANOVA with Tukey multiple comparison test considering p<0.05*, 0.01**, 0.001*** and for (C) Fisher’s exact contingency test (8≤n≤19, N≥3) considering p<0.05*, 0.01**, 0.001***.

The *let-7* miRNA and its downstream target *lin-41*areknown to regulateaxonregeneration pathway and fusion phenomena (Zou et al., 2013, Basu et al., 2017, Wang et al., 2018). Loss of function mutants of *let-7*and *lin-41*showed dendrite regrowth and fusion comparable to that of wild type at24h post-dendrotomy(Figure 4A-C) refuting their role in dendrite regeneration.

PTEN/AKT pathway was previously implicated to play an important role in theregeneration of both axons and dendrites(Huang et al., 2019, Wang et al., 2020). The territory length and fusion phenomenaat 24h post-dendrotomywasnot affected in *akt-1* mutant(Figure3 A-C) suggesting dendrite regeneration in PVD neurons is independent of *akt-1*.

ThePhosphatidylserine (PS) exposure pathway has emerged as an important injury sensing mechanism during axonal injury and dendrite degeneration (Hisamoto et al., 2018, Sapar et al., 2018). Upon injury, the PS signal activates axon regeneration mechanisms such as DLK/MLK p38 MAPK pathway(Pastuhov et al., 2016)or fusogen related repair pathway(Neumann et al., 2015).The PS signal involves exposure of PS to the outer leaflet membrane of injured neuron through the ABC transporter CED-7 and further activation of the downstream effectors such as CED-2/CED-5/CED-12 GEF complex and CED-10 GTPase. This signal subsequently activates p38 cascade involving MLK-1(Pastuhov et al., 2016). We did not see any effect in dendrite regeneration parameters in *ced-7*, *psr-1* and *ced-12* mutants (Figure 4A-C). However, loss of *ced-10* showed a drastic impact on dendrite regeneration including both regrowth and fusion phenomena (Figure 4A-C). In *ced-10* mutant, a large gap is seen at 24h post-dendrotomy since the regrowing branches fail to reach the distal end of the primary dendrites(red dotted line, Figure 4A). The fusion events were also drastically reduced (Figure 4A & C). This indicated that although the PS signal may not be required for dendrite regrowth, the RAC-1 GTPase might have a novel role in the injury response.

The characterization of different axon regeneration pathways in our dendrite regeneration assay clearly indicated that molecular pathways defining dendrite regeneration in PVD neurons have less overlap with the known axon regeneration molecules. But the strong reduction of dendrite regeneration in *ced-10* mutantraised interesting questions to explore further.

### CED-10 RAC GTPaseis required in neuron for dendrite regeneration

Among the candidate genes tested in our dendrite regeneration assay, the*ced-10* mutant showed a strong reduction in dendrite regeneration (Figure 4A-C). Therefore, we investigated the requirement of CED-10 in dendrite injury response in detail.CED-10 is a RAC GTPase involved in regulating cytoskeleton in various morphogenesis processes (Lundquist et al., 2001, Kuhn et al., 1998).The RAC family GTPases have been linked to growth cone navigation during axon development (Quinn et al., 2008). Therefore, we checked whether *ced-10* mutant causes any developmental phenotype in PVD neuron. The length of the axon in PVD neuron remained unaffected in *ced-10* mutant (Figure S3A-B). Dendrites also seemed normal in *ced-10(0)*(Figure S3A). The *ced-10* mutant did not affect axon regrowth parameters also in PVD neuron (Figure S3C-E). To understand the requirement of CED-10 in the initiation of dendrite regeneration, we checked at early timepoints after dendrotomy(Figure 5A). The number of filopodia like structures (arrowheads, Figure 5A)and territory covered seemed to have decreased in*ced-10(0)* as compared to the wildtype at 6h post-dendrotomy (Figure 5A-C).Conversely, when an activated form of CED-10 (G12V) is expressed in the PVD in wildtype background, we found that the number of regrowing branches from proximal dendrite increased at an early timepoint (6h post dendrotomy)(Figure 5A-B). Since the higher concentration lines lead to theformation ectopic branches around the cell body region without even performing dendrotomy, we selected a low concentration line that had very milder defect for our dendrotomy experiment (Figure S3H-I). Similarly, the territory coverage lengthis also increased due to CED-10 activation (Figure 5C). Another gene that codes for RAC GTPase is*mig-2,*which collaborates with CED-10 during development (Lundquist et al., 2001, Kishore and Sundaram, 2002)(Figure S3A). Although, the loss of *mig-2*affected the development of PVD axon (Figure S3A-B), the primary major dendrite regrowth and self-fusion phenomena were unaffected in this mutant (Figure S3F-G).This indicated that developmental impairment of axon would not necessarilyaffect the dendrite regeneration process. This also indicated a specific requirement of CED-10 in the dendrite regeneration of PVD neuron.

**Figure 5.**
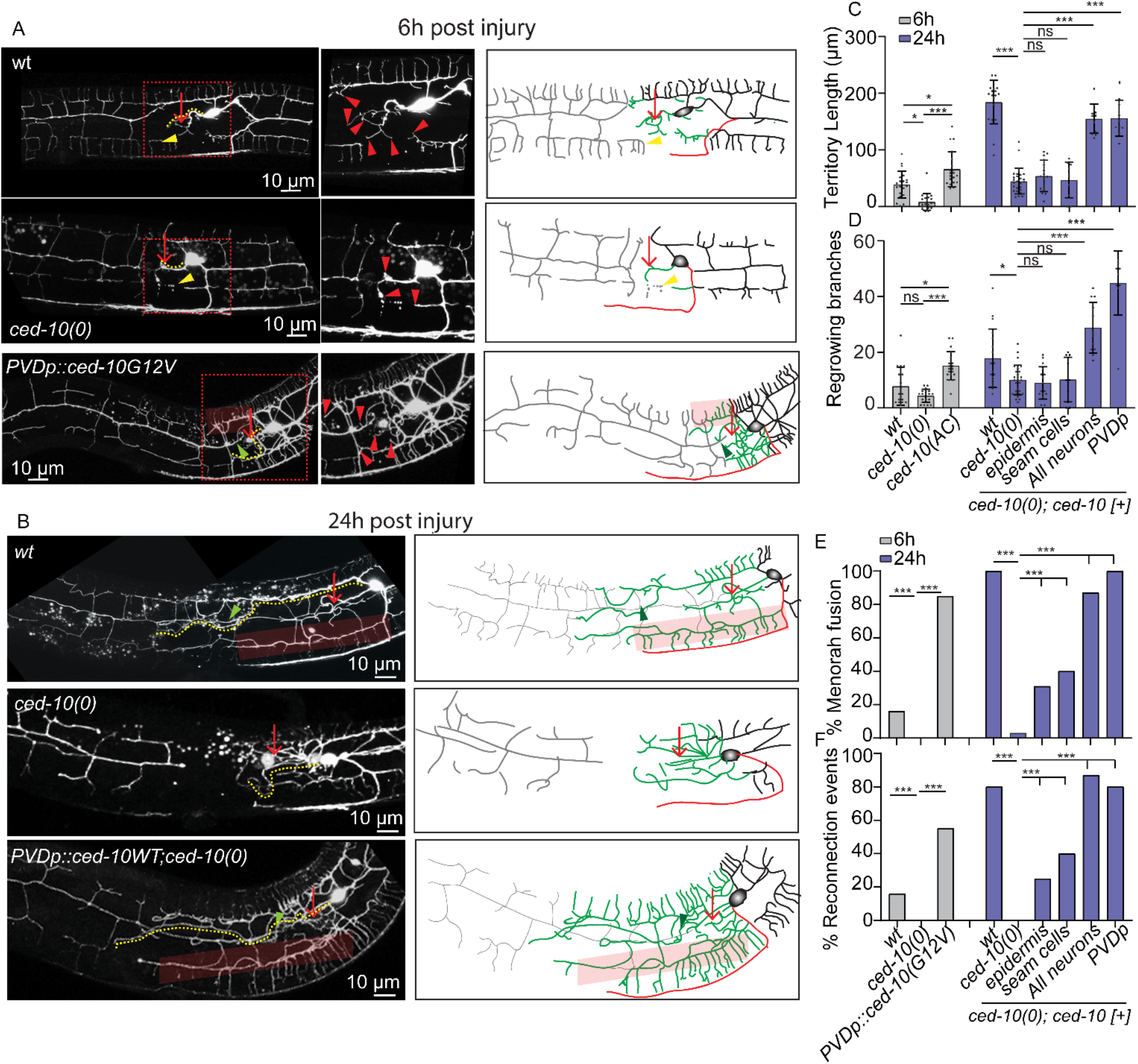
CED-10 is required in neuron for dendrite regeneration. (A) Confocal images of wild type,*ced-10(0)* and *PVDp∷CED-10G12V*(Activated CED-10)at 6h post-dendrotomy along with their schematics indicating site of dendritic injury (red arrow), regrowing dendrites (green), distal part (grey), territory length(yellow dotted lines), reconnection phenomena (green arrowheads) and menorah-menorah fusion (fadedred boxes). Red arrowheads represent the filopodia like structure at the tip of the cut dendrite, yellow arrowheads represent floating degenerating menorah.(B) Confocal images of wildtype, *ced-10(0)* and *PVDp∷ced-10(WT);ced-10(0)* at 24h after dendrotomy along with their schematics indicating site of dendritic injury (red arrow), regrowing dendrites (green), distal part (grey), fusion like phenomena (green arrowheads) and menorah-menorah fusion (fadedred boxes). (C-F)) The quantification of territory length (C), number of regrowing branches (D), the percentage reconnection events (E), and percentage menorah-menorah fusion (F) in Wild Type,*ced-10(0) and PVDp∷ced-10G12V* at6h post-dendrotomy.For the 24h post-dendrotomy time point, thewt, *ced-10(0), pdpy-7∷ced-10(WT)*(epidermis)*;ced-10(0), pgrd-10∷ced-10(WT)*(seam cells)*;ced-10(0),prgef∷ced-10(WT)*(All neurons);*ced-10(0)* and *pser2prom3∷ced-10(WT)*(PVDp);*ced-10(0)* backgrounds were used.For (C_D), (10≤n≤26,N≥3).Statistics, One-way ANOVA with Tukey multiple comparisons method considering p<0.05*, 0.01**, 0.001***. For (E-F) ,(10≤n≤26,N≥3), Statistics,Fisher’s exact test taking p<0.05*, 0.01**, 0.001***.

To check the tissue-specific requirement *ced-10* gene in dendrite regeneration, we expressed the wild type copy of CED-10 under various promoters. We found that when *ced-10* was expressed under pan-neuronal (*prgef)*or PVD-specific promoter *pser2prom3,*the territory length, branch number, % menorah-menorah fusion and reconnection events were completely rescued in *ced-10* mutant background (Figure 5B-C).Surprisingly, when *ced-10* was expressed under the epidermal promoter*pdpy-7*and seam cell promoter *pgrd-10*,we saw a significant rescue of both thereconnection and menorah-menorah fusion, although the territory length and branching were not rescued in this background (Figure 5C-F). This infers that ced-10 RAC GTPase is working cell–autonomously for dendrite regeneration but may also facilitate dendrite regeneration cell-non autonomously by working in nearby epidermal cells.

### TIAM-1 GEF acts upstream of CED-10 in dendrite regeneration

To understand the molecular mechanism by which CED-10 GTPase controls dendrite regeneration in PVD neuron, we speculated that CED-10 could be activated by the upstream factors after dendrotomy. TheRAC GTPases get activated upon the removal of the GDP from their GTP binding domain. This is facilitated by the enzymatic activity of Guanine Exchange Factors (GEFs) (Reiner and Lundquist, 2018). There are some known GEFs for CED-10 such as UNC-73(Trio), TIAM-1(RhoGEF) and CED-12(ELMO1), which contain the RAC binding sites,DH(Dbl homology) and PH(Pleckstrinhomology) domains(Zheng et al., 2016). To identify the relevant GEF of CED-10 in dendrite regeneration, we have performed dendrotomy in the mutants for these GEFs. Although the axons are predominantly missing in the *unc-73(0)* (Figure S4A-B), the dendrite regeneration was unaffected (Figure 6A-C). Similarly,the *ced-12* mutant did not affect any parameters of dendrite regeneration (Figure 6A-C).However, both the territory lengthas well as menorah-menorah fusion events were greatly reduced in the absence of *tiam-1* (Figure 6A-C). The phenotype was very similar to what was seen in the*ced-10* mutant. These phenotypes were completely rescued when the wildtype copy of *tiam-1* is expressed in the PVD neuron under *ser2prom3*(PVD specific) promoter (Figure 6A-D). To test whether CED-10 activation is limiting in *tiam-1*mutant background,we expressed the constitutively activated formof *ced-10* in PVD neuron in the *tiam-1(0)*.The activated form of CED-10 can bypass the requirement of TIAM-1 in both territory extent as well as fusion phenomenain dendrite regeneration (Figure 6B-C).Thus, the RhoGEFTIAM-1acts upstream of CED-10 GTPase for dendritic regeneration.

**Figure 6.**
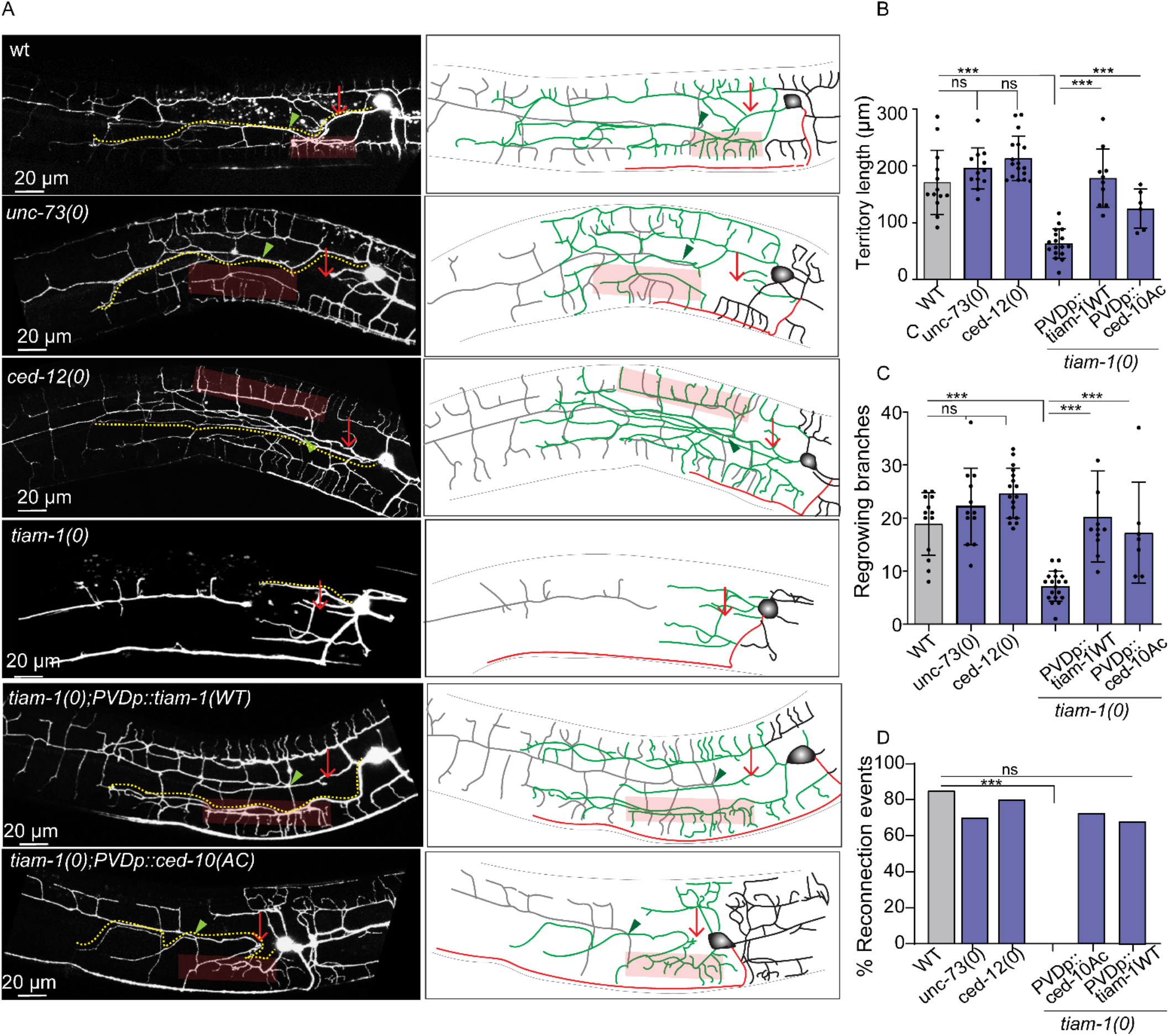
TIAM-1 acts upstream to CED-10 for dendrite regeneration. (A) Confocal images of dendrotomized PVD neuron along with their schematics on the right of *wt, unc-73(0), ced-12(0), tiam-1(0)*,*pser2prom3∷tiam-1WT 5ng; tiam-1(0),* and*pser2prom3∷ced-10(G12V) 5ng;tiam-1(0)*at 24h after injury. The regrowing dendrites are represented as green in color, axon as red and distal part of dendrite as grey in color. green arrowheads represent fusion like phenomena, faded red rectangular boxes indicate menorah-menorah fusion and red arrow marks the site of injury. (B) Quantification of territory length, (C) regrowing branches were measured in *wt, unc-73(0), ced-12(0); tiam-1(0), pser2prom3∷tiam-1(WT);tiam-1(0)*and*pser2prom3∷ced-10(G12V) 5ng;tiam-1(0)* (6≤n≤17,N≥3) and compared using one-way ANOVA by Tukey test considering p<0.05*, 0.01**, 0.001***. (D)The percent of PVD neurons showing reconnection events was measured for*wt, unc-73(0), ced-12(0), tiam-1(0)*, *pser2prom3∷tiam-1WT; tiam-1(0)*and *pser2prom3∷ced-10(G12V);tiam-1(0),* at 24h after dendrotomy (6≤n≤17,N≥3) and compared using two-tailed Fisher’s exact contingency test taking p<0.05*, 0.01**, 0.001***.

## Discussion

In this report, wepresented a detailed analysis of the dendrite and axon regeneration in PVDneuron. Using this system, we could compare the roles of axon regeneration machineries in both dendrite and axon regeneration in the same neuron. Our study revealed a novel function of CED-10 RAC GTPase and TIAM-1 GEF in the dendrite regeneration. TIAM-1/CED-10 cascade is required cell-autonomously in PVD for the initiation of dendrite regrowth and subsequently for branching. Additionally, CED-10 is required in the epidermal cell for regenerative self-fusion events in the same neuron (Figure 7). This expanded our basic understanding of the mechanism of dendrite regeneration.

### PVD neuron as dendrite regeneration model

Dendrite regeneration is poorly studied as compared to the axon regeneration. Recent study using the *da* neuronsin *Drosophila* have shed some light on the mechanism of dendrite regeneration following laser assisted surgery (Song et al., 2012, Stone et al., 2014, Thompson-Peer et al., 2016). Both intracellular as well extracellular machineries control the dendrite regrowth in the *da* neurons(Song et al., 2012, Kitatani et al., 2020, Nye et al., 2020, DeVault et al., 2018).However, the signaling mechanism and downstream effectors that lead to dendrite regeneration is unclear. The dendrites of PVD neuron in *C. elegans* have a stereotypic and an elaborate structure(Inberg et al., 2019). Laser surgery on PVD dendrites leads to self-fusion between the proximal and distal dendrites (Oren-Suissa et al., 2017). The fusion events during dendrite regeneration are driven by fusogenic activity of AFF-1 (Kravtsov et al., 2017). Experiments described in this work allowed addressing the mechanism of dendrite regrowth and branching along withthe fusion process. A comprehensive analysis of the requirement of axon regeneration pathways in dendrite regeneration clearly indicated that the regeneration response to dendrotomy in PVD neuron is largely independent of axon injury response pathways including DLK-1. This is consistent with the finding using *da* neuron in fly (Stone et al., 2014).Our findings show that dendrite regeneration in PVD neuron involves regrowth, branching, and fusion events between the distal and proximal primary dendrites. This is also seen in case of the axon of mechanosensoryPLM neuron that axotomy leads to both regrowth and fusion phenomenon (Ghosh-Roy et al., 2010, Neumann et al., 2011). The finding that that *ced-10* mutant affects both regrowth and fusion events indicates that this pathway is involved in the early response to dendrotomy. Loss of *tiam-1*, the GTP exchange factor for CED-10 GTPase also show similar perturbation in regrowth and fusion events following dendrotomy. This indicated that this conserved RAC GTPase and its upstream activatorTIAM-1 orchestrate the early events after dendrite injury.

### Role of RAC GTPase in dendrite regeneration

The RAC GTPases have been known to play a broader role in developmental processes involving cell debris engulfment (Ellis et al., 1991, Reddien and Horvitz, 2000), migration, axon guidance (Lundquist et al., 2001).The RAC GTPases control these developmental processes through various downstream effectors in actin and microtubule cytoskeleton (Struckhoff and Lundquist, 2003, Quinn et al., 2008, Saenz-Narciso et al., 2016). Previous finding that *ced-10* mutant partially reduces axon regrowth in *C. elegans* motor neuron (Pastuhov et al., 2016)prompted us to test its role in dendrite regeneration. It was seen that *ced-10* acts genetically downstream to Phosphatidylserine (PS) signal to activate DLK/MLK MAP Kinase pathway in regeneration (Pastuhov et al., 2016). However, we did not observe any effect on dendrite regeneration parameters in the mutants affecting either PS or in *dlk-1/mlk-1* mutants ruling out this genetic pathway in the injury response to dendrotomy. However,the cell-autonomous role of TIAM-1 and CED-10 in dendrite regeneration is novel. The RAC GTPases are well-known for their role in F-actin dynamics (Skau and Waterman, 2015, Dominguez and Holmes, 2011), and actin dynamics is a major player in dendritic remodeling during neuronal plasticity (Hotulainen and Hoogenraad, 2010, Chazeau et al., 2014). It might be possible that CED-10 GTPase induces optimal F-actin dynamics suited for regrowth and branching that is observed after dendrite injury.

Our finding that CED-10 plays a cell non-autonomous function in the surrounding epithelial cells for the menorah-menorah fusion events during regeneration is very intriguing. It was seen that for the fusion to take place, AFF-1 fusogen was delivered from the surrounding seam cells, which are of epithelial origin. It is possible that CED-10 initiates the epidermal response to dendrotomy, which might lead to the release of vesicles containing AFF-1 from epidermal cells. Epidermal cells are known for responding to dendrite injury. In case of *da* neuron in *Drosophila*, the PS pathway controls the dendrotomy induced engulfment of degenerated distal dendrites(Han et al., 2014, Sapar et al., 2018).

## Materials and Methods

### *C. elegans* strains and genetics

The *C. elegans* strains were grown and maintained at 20°C in OP50 bacterial lawn seeded in Nematode Growth Medium (NGM) plates(Brenner, 1974). The loss of function mutation is represented as (0) for example loss of function allele of *dlk-1, tm4024* would be represented as *dlk-1(0).*The mutants that were used in this study are mostly loss of function by deletion or substitution unless otherwise mentioned (Table S1). These mutants were taken from Caenorhabditis Genetics Centre (CGC) and genotyped using their respective genotyping primers (Table S2).

### Molecular cloning and creating transgenes

Destination vector with PVD neuron specific promoter, *pser2*[4.1kb]∷Gateway [pNBRGWY99] was made by infusion cloning (Takara). *ser2* Promoter region was amplified from the fosmid WRM0623bG06 using the primers: 5’-ccatgattacgccaagtaaaagtttagtaaattaactgc-3’ and 5’-tggccaatcccggggtatgtgttgtgatgtcac-3’ and GWY vector backbone was amplified from pCZGY553 using 5’-ccccgggattggcca-3’ and 5’-ttggcgtaatcatgg-3’.

For pan-neuronal and epidermal rescue of *ced-10*,*prgef*∷Gateway [pCZGY66] and *pdpy-7*∷GWY [pNBRGWY44], respectively, were recombined with the entry clone of *ced-10 WT* [pNBRGWY88] (Puri et al., 2021)using the LR recombination(Invitrogen).

For seam cell specific rescue, the promoter *pgrd-10* was amplified from fosmid WRM0612bC07 using 5’-ccatgattacgccaatcgtcatc-3’ and 5’-tggccaatcccggggtttttaga-3’ and cloned with Gateway backbone amplified using 5’-ccccgggattggcca-3’ and 5’-ttggcgtaatcatgg-3’ to make *pgrd-10∷GW* [pNBRGWY151] using infusion cloning(Takara). It was then recombined with *ced-10* WT[pNBRGWY88] using LR recombination (Invitrogen).

For PVD specific rescue of wildtype and constitutively active *ced-10*, pNBRGWY99 was recombined with *ced-10 WT* [pNBRGWY88] and *ced-10 constitutively active (G12V)* [pNBRGWY89] plasmids(Puri et al., 2021), respectively, using the LR recombination(Invitrogen).

*tiam-1* cDNA was cloned intopNBRGWY99 by infusion cloning for PVD specific expression of *tiam-1*. Primers used to amplify tiam-1 cDNA from total cDNA were 5’-tccgaattcgcccttatgggctcacgcctctca-3’, and 5’-aaggaacatcgaaattcaaaatagcagctttcttgtaca-3’.The primers used to amplify vector backbone are 5’-cagctttcttgtaca-3’ and 5’-aagggcgaattcgga-3’.

The clones made using various molecular techniques were then injected in the gonad of young adult worms along with co-injection markers such as *pttx-3∷RFP* or *pmyo-2∷mCherry* and the F1 progeny were then isolated and checked for the formation of transgenic lines. Each transgene was checked for two high transmission lines. List of transgenes used in this study are listed in table S3.

### Laser system, dendrotomy and axotomy

Dendrotomy and axotomy experiments were conducted at the L4 stage of *C. elegans* using the Bruker® ULTIMA system with SpectraPhysics®Two-photon femtosecondlaser which is a tunable Infra-red (690-1040 nm) laser having automated dispersioncompensation(MaiTai with DeepSee)(Basu et al., 2017).The output of laser was controlled using Conopticspockel cells with superior temporal resolution (~μs) and galvanometer having range of 6mm and 3mm were used for imaging and severing, respectively. 920nm and 720nm laser (pulse width~80fps, irradiation pulse width: 20ms, point spread function (PSF) xy~400nm and z~1.5μm) were used simultaneously, for the visualization and severing of PVD, respectively. Worm containing slide was imaged using 60X/0.9NA water objective (Olympus®) at a pixel resolution of 0.29μm x 0.29μm. Worms were immobilized using either Levamisole hydrochloride (10mM) (Sigma L0380000) as a paralyzing drug agent or with polystyrene beads (Polysciences 00876-15) of diameter 0.1 μm as friction enhancing agent on 5% agarose pads mountedwith Corning cover glass (cat No. 2855-25). Nikon Ti-2 microscope equipped withfemtosecond Ultra-violet laser (Micropoint laser®, 337nm, 3ns Pulse, Max Pulse enery-200μJ, Max Average Power-4mW) system was also used to assist the laser injury using using 100x oil immersion with NA 1.40.

PVD dendrites were severed with the first laser shot at the first branch point (~10μm away from cell body)followed by one (two shot experiment, Fig 1A) or more (multi shot experiment, FigS1A) consecutive shots with a relative distance of 10-15μm from the previous shot creating a visible gap with no fragments left at the cut site. The axons were severed distally at the segment fasciculating with the ventral nerve cord (VNC) (~50 μm away from cell body). A clean transection was ensured by making two shots 5 μm apart at a distance of 10 μm away from the distal most lateral to ventral transition (LV) point of PVDR and PVDL.

After severing, worms were recovered from the agarose pad using an aspirator with 1X M9 solution onto freshly seeded NGM plates for further observation.

### Imaging

To observe the regenerative response, injured worms were imaged at 3, 24 and 48 hours (h) after injury. The worms were paralyzed and mounted in 10 mM Levamisole hydrochloride (Sigma®) solution on slides containing medium on 5% agarose (Sigma®) pads. The worms were imaged with 63X/1.4NA oil objective of Nikon® A1R confocal system at a voxel resolution of 0.41μm x 0.41μm x 1μm and tile imaging module using imaging lasers 488nm(GFP), 543nm(mCherry/RFP) with 1-1.8 AU pinhole at 512×512 pixel resolution files for further analysis.

### Dendrite Regeneration Analysis and Quantification

Dendrite regeneration was quantified on the basis of regrowth and fusion parameters. The territory covered by the regenerated dendrite (figure 1B, yellow dotted line) was estimated using Simple Neurite Tracer plugin in Fiji-ImageJ® tracking the longest regrowing dendrite from the cell body to the tip of the dendrite. ‘Total number of branches’ were then also calculated using the same plugin from the point of emergence (cell body or dendrite) to the termination of every branch.

PVD dendrites have a capability to fuse after injury (Kravtsov et al., 2017, Oren-Suissa et al., 2017). We also observed reconnection between the regenerating proximal primary dendrite with distal primary dendrite or distal menorah (figure 1B, green arrow) which were evaluated as reconnection events, the population of worms showing this phenotype was calculated and represented as a fraction. Long menorahhaving more than one secondary dendrite connected to it (figure 1B, faded red rectangular boxes), was classified as menorah-menorah fusion and percentage of PVD neuron showing menorah-menorah fusion was quantified. Following injury, distal dendrite may show hallmarks of Wallerian degeneration (Kerschensteiner et al., 2005, Waller, 1851), which was estimated as the percentage of PVD neurons showing distal part degeneration (figure 1C, grey labelled dotted neurites).

### Axon Regeneration Analysis and Quantification

Analysis of the axon regeneration was carried out by visual observation and quantitative information were obtained using various image analysis modules of ImageJ®. Neurites were traced and their lengths were quantified using the Simple Neurite Tracer plugin of ImageJ®. Volumetric visualization facilitated obtaining a qualitative estimate and descriptors of regenerated neurites based on the mCHERRY∷RAB-3 localization. Regenerative growth was classified into the neurite growth from the cut point of the axon, conversion of the adjacent tertiary dendrites to axon like identity as per the localization of mCHERRY∷RAB-3punctae, ectopic branches from the cell body, proximal axon, or converted neurites.

Quants obtained from ImageJ® for both dendritic and axonal regeneration were further analyzed by Excel and Graphpad® to get statistical information.

### Statistical analysis

Statistical analyses were compassed using Graphpad Prism software (Prism 8 V8.2.1(441)). Two samples were analyzed using unpaired two-tailed t test. The statistical analysis of multiple samples was performed using One way-ANOVA with Tukey multiple comparisons test. The data that was used for ANOVA analysis was naturally occurring data having normal distribution spread which was not further processed. To compare the population data, the fraction values with respect to each sample were calculated and compared using the two-tailed Chi square Fisher’s exact contingency test was performed. The biological replicates for each experiment were done at least three times. The significance considered for all statistical experiments are p<0.05*, 0.01**, 0.001***.

## Acknowledgments

We thank Yuji Kohara for cDNAs. We thank National BioResource Project (NBRP), Japan, and Caenorhabditis Genetics Center (CGC) for strains. CGC is supported by the NIH Office of Research Infrastructure Programs (P40 OD010440). We thank Sandhya Koushika, and Cori Bargmann for the help with strains. We thank Erik A. Lundquist for providing reagents to manipulate small GTPases. This work is supported by the NBRC core fund from the Department of Biotechnology, DBT/Wellcome Trust India Alliance (Grant # IA/I/13/1/500874 to A.G.-R. and Grant # IA/E/18/1/504331 to S.D.).

## Author Contributions

H. Kaur Brar, Swagata Dey, and A. Ghosh-Roy designed experiments. H. Kaur Brar, Swagata Dey, and D. Pandey performed experiments and analyzed data. H. Kaur Brar, Swagata Dey, and A. Ghosh-Roy wrote the manuscript. Swagata Dey developed axotomy protocol, performed other axotomy-related experiments and analysis. H. Kaur Brar developed the dendrotomy protocol, performed dendrotomy-related experiments and analysis. D. Pande did dendrotomy-related experiments analysis. S. Bhardwaj, and H. Kaur Brar did crosses, designed constructs and performed cloning. P. Singh designed constructs and performed cloning. Shirshendu Dey maintained and optimized the lasers in the 2-photon microscope.

