## Supplementary Data for "Dendrite regeneration in *C. elegans*is controlled by the RAC GTPase CED-10 and the RhoGEF TIAM-1"

### List of supplementary figures

**Figure S1:** Regeneration response of primary dendrites of PVD, Related to Figure 1**(3-4)**

**Figure S2:** The dendrite regeneration does not require DLK/MLK pathway, Related to Figure 2 **(5-6)**

**Figure S3:** CED-10 is required for dendrite regeneration, Related to figure 5 **(7-8)**

**Figure S4:** Developmental imaging PVD neuron in Rho/RAC-GEF mutants, related to Figure 6 **(9)**

### List of Tables

**Table S1:** List of *C. elegans* strains used in this paper **(10)**

**Table S2:** List of strains carrying extrachromosomal transgenes used in this paper **(11-12)**

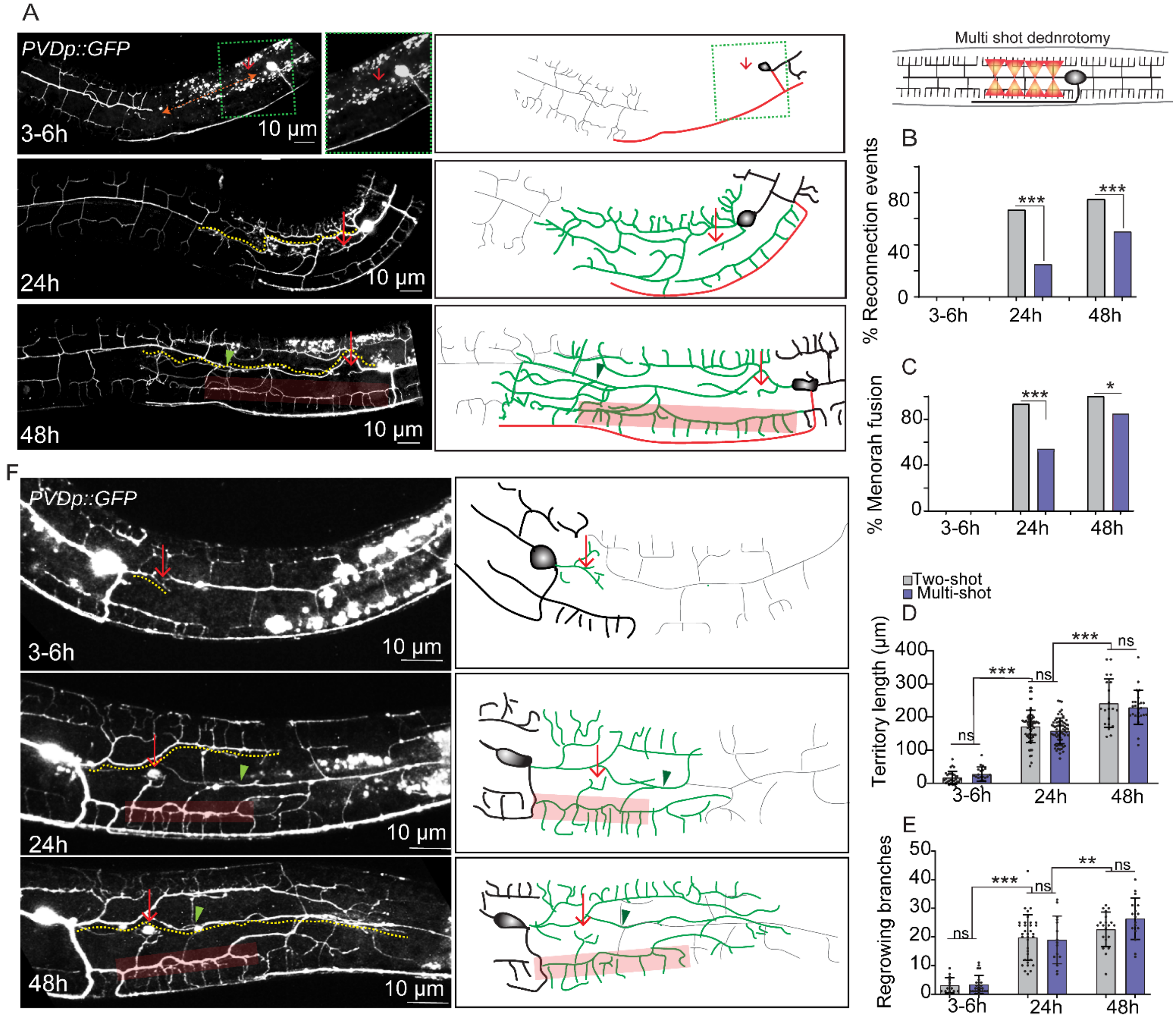


**Figure S1. Regeneration response of primary dendrites of PVD, Related to Figure 1**
(A) The confocal images and illustrations (right) of regeneration phenomena of primary major dendrites of PVD at 3h, 24h and 48h after dendrotomy using four laser shots. The experiment was performed in worms expressing *wdIs52* (*pF49H12.4::GFP*) reporter. The large gap created due to this multi-shot dendrotomy at 3h indicated with orange dotted line with double-arrowheads (topmost panel). The semi-transparent red box and green arrowhead highlight the menorah-menorah fusion event and reconnection phenomenon, respectively. In the illustration (right), the green neurites indicate regrowing dendrites, grey indicates remnants of distal part of dendrite and the axon is marked in red.

(B-C) The percentage occurrence of reconnection events ($20 \leq n \leq 60, N \geq 3$), and menorah-menorah fusion ($20 \leq n \leq 60, N \geq 3$). The Fisher exact two-tailed test was done for statistical analysis. ns, non-significant, $p<0.05$*, $p<0.01$**, 0.001***.
(D-E) Quantification of territory length ($20 \leq n \leq 70, N \geq 3$)(D), and the total number of branches ($14 \leq n \leq 34, N \geq 3$)(E) at 3h and 24h post-dendrotomy during single and multi-shots of each timepoint compared within and across timepoints using one-way ANOVA with Tukey multiple comparisons method. Data is represented as a column scatter and histogram of Mean$\pm$S.D. $p<0.05^{*}$, $0.01^{**}$, $0.001^{***}$, ns (non-significant, $p>0.05$).
(F) Confocal images with schematics showing the regeneration events at 3, 24, and 48 h following the injury on primary minor dendrites.

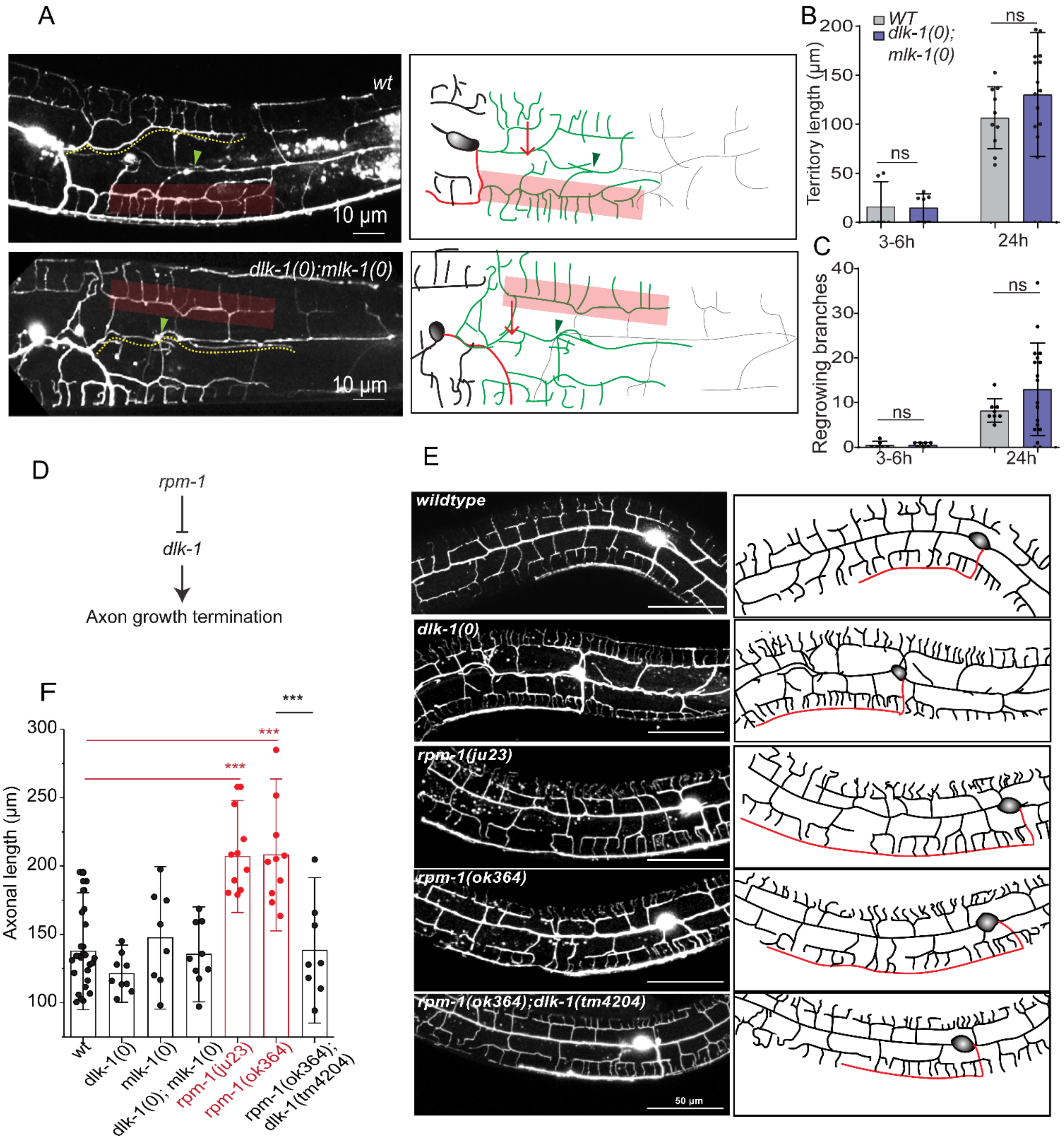


**Figure S2. The dendrite regeneration does not require DLK/MLK pathway, Related to figure 2**

(A) Confocal images of the regeneration events of minor dendrites in WTand *dlk-1(0);mlk-1(0)* backgrounds at 24h post-dendrotomy. The schematics representing site of dendritic injury (red arrow), regenerated dendrite (green), distal part (grey), reconnection phenomena (green arrowheads) and menorah-menorah fusion (semi-transparent red boxes).Territory length is represented as yellow dotted lines in the confocal image(B-C) The territory length and the number of regrowing branches, and ($7 \leq n \leq 19$ $N=3$) (C) at 3-6h and 24h post-dendrotomy. ($7 \leq n \leq 19$ $N=3$). Statistics, ANOVA with Tukey multiple comparisons method taking $p<0.05$*, 0.01**, 0.001***. (D) The genetic pathway of*rpm-1* pathway controlling axon growth termination. (E) Representative confocal images showing the developmental phenotype of PVD in various mutants of *rpm-1* pathway. In the schematics, the axon in shown in red. Please note that in rpm-1 mutants an overshooting of axon phenotype is noticed. (F) The quantification of axonal length of PVD neurons in the wild-type, *dlk-1(0)*, *mlk-1(0)*, *dlk-1(0); mlk-1(0), rpm-1(ju23), rpm-1(ok364) and rpm-1(ok364);dlk-1(tm4024)* mutants at L4 stage($8 \leq n \leq 27$, $N \geq 3$). The statistical comparison was done using one-way ANOVA Bonferroni test considering $p<0.05$*, 0.01**, 0.001***.

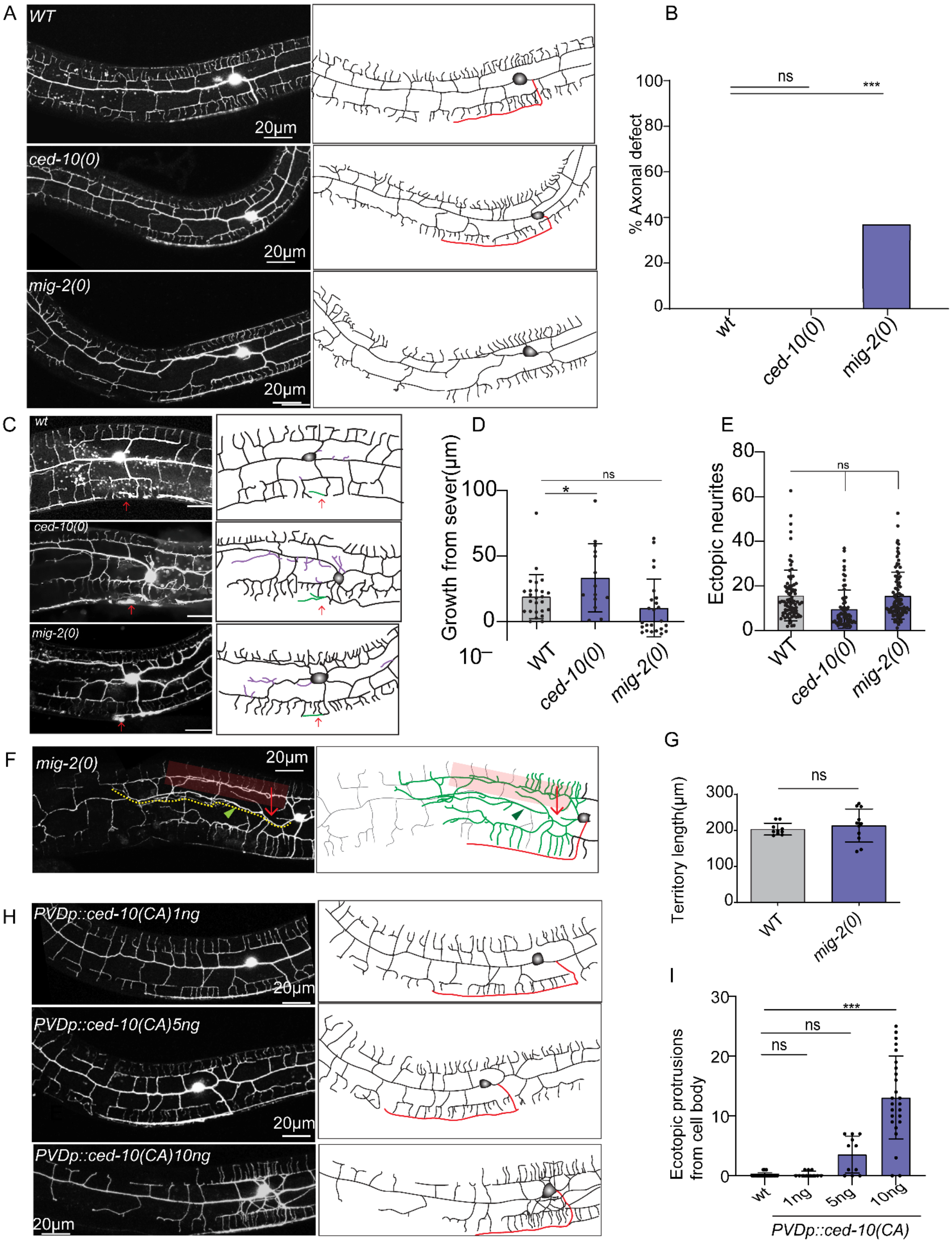

A
WT
20µm
ced-10(0)
20µm
mig-2(0)
20µm
B
% Axonal defect
100
80
60
40
20
0
ns
***
wt
ced-10(0)
mig-2(0)
C
wt
ced-10(0)
mig-2(0)
D
Growth from sever(µm)
100
50
0
10
*
ns
WT
ced-10(0)
mig-2(0)
E
Ectopic neurites
80
60
40
20
0
ns
WT
ced-10(0)
mig-2(0)
F
mig-2(0)
20µm
G
Territory length(µm)
300
200
100
0
ns
WT
mig-2(0)
H
PVDp::ced-10(CA)1ng
20µm
PVDp::ced-10(CA)5ng
20µm
PVDp::ced-10(CA)10ng
20µm
I
Ecotopic protrusions from cell body
30
20
10
0
***
ns
ns
wt
1ng
5ng
10ng
PVDp::ced-10(CA)

**Figure S3. CED-10 is required for dendrite regeneration, related to Figure 5.**

(A) Confocal imaging of *wt, ced-10(0),* and *mig-2(0)* is shown along with its schematics(right) indicating the axon in red and its quantification of (B) axonal defect at L4 stage on population basis is represented as percentage defect ($12 \geq n \geq 10$) and compared using Fisher's exact two-tailed test considering $p<0.05$*, 0.01**, 0.001***.
(C) Confocal images of *wt*, *ced-10(0)* and *mig-2(0)* at 24h after axotomy at around 50 μm away from cell body along with their schematics indicating site of axonal injury with red arrow, regrowing axon in green from severed end and purple representing ectopic neurites.
(D-E) Quantification of axon regeneration as growth from the severed end (D) and Length of ectopic neurites(E), of *wt, ced-10(0)* and *mig-2(0)* at 24 h after axotomy ($25 \geq n \geq 14$, $N \geq 3$) and compared using one-way ANOVA Tukey method taking $p<0.05$*, 0.01**, 0.001***.
(F) Confocal images of *mig-2(0)* at 24h after dendrotomy is shown along with its schematics indicating regrowing dendrites as green color, distal part as grey color, fusion like phenomenon with green arrowheads, menorah-menorah fusion with faint red rectangular boxes and red arrow marking the site of axotomy.
(G) Dendrite regeneration parameter i.e. territory length of *wt*and *mig-2(0)* at 24h after dendrotomy ($10 \leq n \leq 11$, $N \geq 3$) is plotted and compared using two-tailed t test considering $p<0.05$*, 0.01**, 0.001***.
(H) Confocal images of *pser2prom3::ced-10(G12V) 1ng/μl, pser2prom3::ced-10(G12V) 5ng/μl,* and *pser2prom3::ced-10(G12V) 10ng/μl* at L4 stage is shown along with their schematics to represent dendrites (black) and axon(red).
(I) Quantification of number of ectopic neurites emerging out of cell body or adjacent dendrites in *wt*, *pser2prom3::ced-10(G12V)1ng*, *pser2prom3::ced-10(G12V)5ng,* and *pser2prom3::ced-10(G12V)10ng* at L4 stage ($11 \leq n \leq 20$, $N=3$) and compared using one way ANOVA Tukey test taking $p<0.05$*, 0.01**, 0.001***.

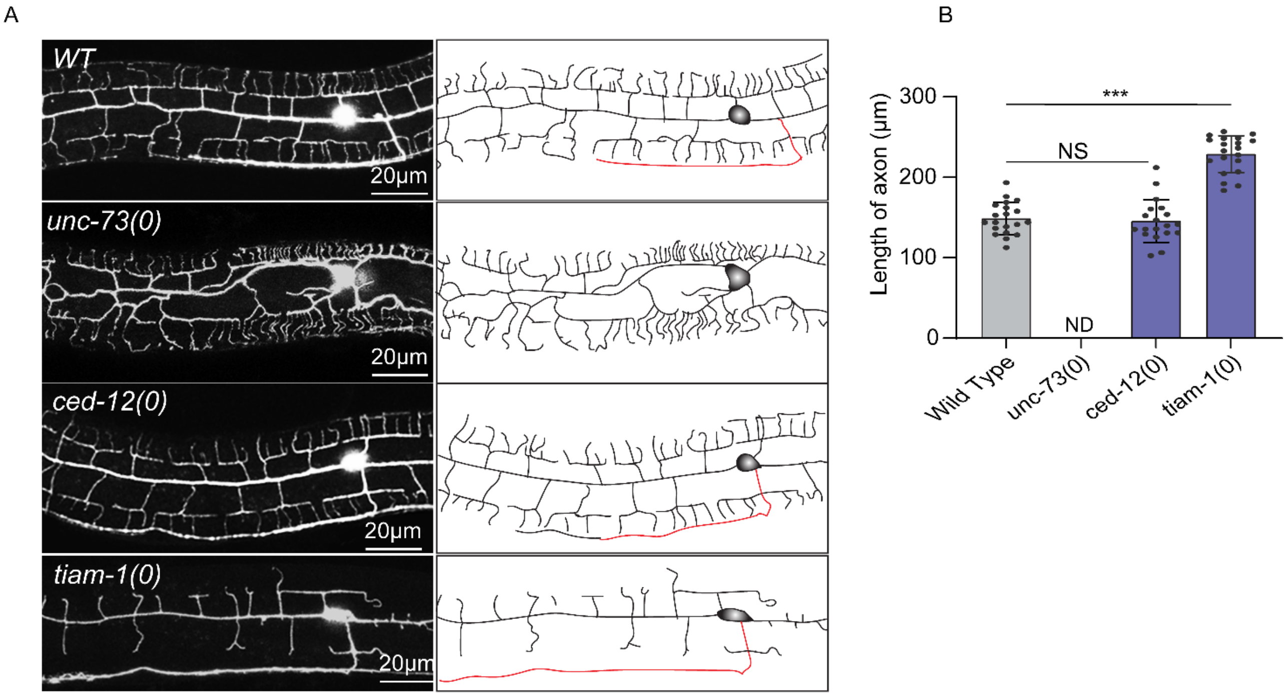


**Figure S4: Developmental imaging PVD neuron in Rho/RAC-GEF mutants, related to figure 6**

(A) Confocal images of *wt, unc-73(0), ced-12(0), tiam-1(0)* and *pser2prom3::ced-10(G12V) ;tiam-1(0)* along with their schematics showing the developmental phenotype in PVD neuron at L4 stage.

(B) Length of axon is calculated for *wt, unc-73(0), ced-12(0)* and *tiam-1(0)* (21≥n≥20,N=3) and compared using one-way ANOVA with Tukey method considering p<0.05*, 0.01**, 0.001***.

**Table S1 List of C. elegans strains used in this paper**

| ***C.elegans*strains** | **Source** | **Identifier** |
|---|---|---|
| *WdIs52(F49H12.4::GFP, unc119[+]) X* | CGC | NC1687 |
| *des-2::mCherry::RAB-3, des-2::SAD-1::GFP(kyIs445)* | Cori Bargmann Lab | CX9797 |
| *dlk-1(tm4024)I* | CGC | CZ15956 |
| *mlk-1(ok2471)I* | CGC | RB1908 |
| *rpm-1(ju23)V* | CGC | CZ1234 |
| *rpm-1(ok364)V* | CGC | RB630 |
| *let-7(mg279)X* | CGC | GR1432 |
| *lin-41(ma104)I* | CGC | CT8 |
| *akt-1(mg306)V* | CGC | BQ1 |
| *psr-1(tm469)IV* | CGC | CU1715 |
| *ced-7(n1996)III* | CGC | MT4983 |
| *ced-12(bz187)I* | CGC | ZB547 |
| *pde-4(ce268)II* | CGC | KG744 |
| *egl-19(ad695)IV; muIs32(pmec-7::GFP)II* | CGC | CZ9927 |
| *ced-10(n3246)IV* | CGC | MT9958 |
| *max-2(cy2)II* | CGC | VC1462 |
| *mig-2(ok2273)X* | CGC | RB1769 |
| *tiam-1(ok772)I* | CGC | RB907 |
| *unc-73(rh40)I* | CGC | LE137 |

**Table S2: List of stains carrying extra chromosomal array used in this paper**

| Strain | Transgene | DNA Construct | Genetic Background | DNA concentration |
|---|---|---|---|---|
| NBR775 | *shrEx387* | pNBRGWY136(*prgef::CED-10 WT)* | *ced-10(n3246); WdIS52* | 10ng/µl |
| NBR776 | *shrEx388* | pNBRGWY136(*prgef::CED-10 WT)* | *ced-10(n3246); WdIS52* | 10ng/µl |
| NBR796 | *shrEx399* | pNBRGWY141(*pdpy-7::CED-10WT)* | *ced-10(n3246); WdIS52* | 10ng/µl |
| NBR956 | *shrEx455* | pNBRGWY141(*pdpy-7::CED-10WT)* | *ced-10(n3246); WdIS52* | 1ng/µl |
| NBR957 | *shrEx456* | pNBRGWY141(*pdpy-7::CED-10WT)* | *ced-10(n3246); WdIS52* | 1ng/µl |
| NBR946 | *shrEx446* | pNBRGWY154(*pgrd-10::CED-10*) | *ced-10(n3246); WdIS52* | 5ng/µl |
| NBR947 | *shrEx447* | pNBRGWY154(*pgrd-10::CED-10*) | *ced-10(n3246); WdIS52* | 5ng/µl |
| NBR902 | *shrEx422* | pNBRGWY121(*pser2prom3(4.1kb)::CED-10WT)* | *ced-10(n3246); WdIS52* | 10ng/µl |
| NBR903 | *shrEx423* | pNBRGWY121(*pser2prom3(4.1kb)::CED-10WT)* | *ced-10(n3246); WdIS52* | 10ng/µl |
| NBR904 | *shrEx424* | pNBRGWY130(*pser2prom3(4.1kb)::CED-10(G12V))* | *WdIS52* | 5ng/µl |

| | | | | |
|---|---|---|---|---|
| NBR905 | *shrEx425* | pNBRGWY130(*pser2prom3(4.1kb)::CED-10(G12V))* | *WdIS52* | 5ng/µl |
| NBR906 | *shrEx426* | pNBRGWY130(*pser2prom3(4.1kb)::CED-10(G12V))* | *WdIS52* | 1ng/µl |
| NBR907 | *shrEx427* | pNBRGWY130(*pser2prom3(4.1kb)::CED-10(G12V))* | *WdIS52* | 1ng/µl |
| NBR931 | *shrEx452* | pNBRGWY130(*pser2prom3(4.1kb)::CED-10(G12V))* | *WdIS52* | 10ng/µl |
| NBR932 | *shrEx453* | pNBRGWY130(*pser2prom3(4.1kb)::CED-10(G12V))* | *WdIS52* | 10ng/µl |
| NBR910 | *shrEx428* | pNBRGWY130(*pser2prom3(4.1kb)::CED-10(G12V))* | *tiam-1(0); wdIs52* | 5ng/µl |
| NBR911 | *shrEx429* | pNBRGWY130(*pser2prom3(4.1kb)::CED-10(G12V))* | *tiam-1(0); wdIs52* | 5ng/µl |
| NBR944 | *shrEx444* | PNBR46 (*pser2prom3(4.1kb)::tiam-1cDNA)* | *tiam-1(0);wdIs52* | 5ng/µl |
| NBR945 | *shrEx445* | PNBR46 (*pser2prom3(4.1kb)::tiam-1cDNA)* | *tiam-1(0);wdIs52* | 5ng/µl |
